# Benchmarking phenology with a land surface model: an in- silico experiment for temperate forests

**DOI:** 10.1101/2025.11.20.689458

**Authors:** Midori Yajima, Josua I. Seitz, William J. Matthaeus, Gayathri G. Nair, Luke Daly, Silvia Caldararu

## Abstract

Plant phenology influences biotic interactions and the cycling of water, carbon, and nutrients, ultimately affecting global climate. Accurately quantifying phenological dynamics is therefore essential for ecological forecasting, yet estimates vary widely depending on data sources, smoothing approaches, and date-extraction techniques. For this reason, the selection of an appropriate processing pipeline requires benchmarking under controlled conditions. In this study, we use a Land Surface Model (LSM) to simulate ecosystem dynamics at four temperate forest sites under five lesser studied climate perturbations. The resulting synthetic dataset allows us to control for environmental conditions while comparing commonly used phenological detection methods (relative thresholds at 10%, 20% and 50%, highest rate of change, detection of inflection points in the curvature’s angle, detection of recovery and senescence points). Amongst tested methods, we find general agreement in responses to climate alterations, where inflection and low thresholds (10%) methods provide the closest match to the model’s predicted start of season, with a maximum discrepancy of 4 days. Higher thresholds (20%, 50%) are closer to the model’s predicted end of season, though discrepancies reach up to 10 days. High thresholds and rate-of-change methods also reliably detect the peak of season. We further show that some methods detect phenological responses missed by temperature-only type growing season, supporting the need for more realistic phenological representations in LSMs. Beyond evaluating extraction methods within a common framework, our study expands on the potential of using LSMs as sandboxes for ecological testing rather than solely predictive tools.

## 1 Introduction

Phenology refers to the timing of seasonal events and their biotic and abiotic drivers (Richardson et al., 2013). In extra-tropical regions, seasonality drives different aspects of plant activity, including plant growth, net carbon gain, or occurrence of specific events (e.g. budburst, flowering, leaf greenup, coloration and fall), that commonly define a growing season (Körner et al., 2023). The timing of these events shapes reproductive success, biotic interactions, and ecosystem functioning through water, nutrient and carbon cycling, ultimately affecting the ecosystems’ energy balance (Forrest & Miller-Rushing, 2010; Katal et al., 2022; Silvestro et al., 2025; Chen et al., 2022). This results in feedback between terrestrial ecosystem functioning and the global climate, where the timing of phenological events has direct consequences on the climate system on one side, but also responds to climatic changes (Keenan et al., 2014) with increased temperatures and shifts in precipitation patterns affecting onset and duration of a growing season (Campioli et al., 2024).

Assessing and modelling phenology is therefore central to understand how ecosystems both contribute and respond to environmental conditions (Katal et al., 2022). Traditionally, phenological research investigates data and metrics related to temperature responses of deciduous species in temperate climates, and numerous studies have focused on the effects of warmer springs on advancing the growing season (Hassan et al., 2023; Shi et al., 2025; Stuble et al., 2021), and to a lesser degree on the effects of warmer autumns on delaying leaf off (Calinger & Curtis, 2023). However, globally, many other seasonal patterns exist including dry-wet season dynamics in the tropics (Abernethy et al., 2018) and functional seasonality in boreal evergreen forests (Yang et al., 2020). Even in temperate regions change in environmental conditions beyond temperature can affect seasonality. For example, dry springs characterized by high solar radiation have been shown to lead to earlier greening and increase in productivity (Bastos et al., 2020).

In addition, our understanding of phenology is also limited by the variety of data source and detection methodologies used to derive growing season metrics. Phenological data sources span from traditional point observations of leaf-on and leaf-off timing, high resolution ground measurements of greenness from cameras and productivity from eddy covariance measurements to decades long timeseries of remotely sensed vegetation indexes (Gallinat et al., 2021; Gray & Ewers, 2021; Katal et al., 2022; Reyes-González et al., 2021). From times series of data sources we then generally derive growing season metrics through a variety of mathematical de-noising techniques and transition dates detection methods (Caparros-Santiago et al., 2021; Klosterman et al., 2014; Panwar et al., 2023). This leads to uncertainty as the type of observation measures different ecosystem properties, and the mathematical model and phenological extraction method detects different points in the growing season. Further, while traditional tree phenology is concerned with the time of leaf appearance and disappearance, by either direct observation or calculation from greenness timeseries, the study of ecosystem carbon dynamics focuses more on the seasonality of productivity, frequently measured or modelled as Gross Primary productivity (GPP). While leaf growth and productivity are inherently interlinked, the dependency is non-linear and the phenological impacts could differ (Körner et al., 2023). This not only limits our understanding of vegetated systems’ response to climate change, but also their ecological forecasting, as the evaluation of model performances would be based on non-representative phenological metrics.

The selection of the most appropriate source of data and processing for the system of study should therefore follow benchmarking, to determine whether detected differences in the derived growing season are climate-or methodology-related. This is however difficult to determine from experimental and observational data, where the interaction between climatic drivers and local conditions further complicates the attribution of phenological patterns (Wolkovich et al., 2012). For this reason, in this study we simulate four temperate forest sites with the QUINCY (QUantifying Interactions between terrestrial Nutrient CYcles and the climate system, Thum et al., 2019) Land Surface Model (LSM) to provide an *in silico* experimental setting, i.e. a synthetic representation of key vegetation processes that respond to fully controlled environmental conditions. We then test four commonly used methodologies to extract the start (SOS) and end (EOS) of a growing season on simulated carbon uptake (Gross Primary Productivity, GPP) and canopy dynamics (Leaf Area Index, LAI) under five lesser studied climate scenarios. We subsequently compare the derived growing season metrics against QUINCY’s in-built temperature dependent growing season. This aims to assess how phenological transition dates vary across detection methods accounting for different plant processes, the sensitivity of each method to detect climate changes, as well as how they compare against a theoretical ‘true’ growing season.

## 2 Materials and Methods

### 2.1 Site Description

We selected four sites dominated by deciduous broadleaf forest (DBF) from the PLUMBER2 dataset (Ukkola et al., 2022), which comprises of 170 flux tower sites globally for use in land surface modelling. The selected sites, Hainich (DE-Hai), Fontainebleau-Barbeau (FR-Fon), Harvard Forest (US-Ha1), and Morgan Monroe State Forest (US-MMs), span a range of mean annual temperature (MAT) and precipitation (MAP), reflecting the climatic extent of temperate broadleaf forests in the dataset (Figure B1). We did not consider representativeness of species or soil type, and only meteorological data from the sites were used to drive the model. However we observe similar structural characteristics between sites, with a woody vegetation cover > 60% and dominant vegetation type representative of deciduous broadleaf forests (Berveiller et al., 2005–2014; Knohl et al., 2000–2012; Munger, 1991–2012; Novick & Phillips, 1999–2014). An overview of site characteristics can be found in Table 2.

**Table 1.** Abbreviations used in this study and their description.

| Abbreviation | Term | Description |
| --- | --- | --- |
| DBF | Deciduous<br>Broadleaf Forest | Forest dominated by broad-leaf trees that shed seasonally (also referred to as temperate forest in this study) |
| DER | Derivative Method | First derivative of the vegetation time series |
| DOY | Day of Year | Sequential day number within a year |
| EOS | End of growing<br>Season | Point in time at which vegetation activity begins to decline,<br>marking the transition from active growth to dormancy,<br>senescence, or leaf fall (typically autumn) |
| GDD | Growing Degree<br>Days | Index used to estimate plant development based on temperature<br>accumulation over time (measured in °C days) |
| GPP | Gross Primary<br>Productivity | Carbon uptake of vegetation before accounting for carbon loss<br>through respiration (measured in mol/ day in this study) |
| GU | Gu Method | Intersection between upper and lower boundaries of vegetation<br>time series and its recovery and senescence line |
| INF | Inflection method | Rate of change in the angle of the tangent vector for the<br>vegetation time series |
| LAI | Leaf Area Index | Total area of photosynthetic surface per unit ground area |
| LSM | Land Surface<br>Model | Mathematical representations of physical and biogeochemical<br>processes governing the exchange of energy, water, and nutrients<br>between land and the atmosphere |
| MAP | Mean Annual<br>Precipitation | Average precipitation over the course of a year |
| MAT | Mean Annual<br>Temperature | Average air temperature over the course of a year |
| PFT | Plan Functional<br>Type | Classification of vegetation types grouping plants species based<br>on shared functional traits (e.g. physiology, phenology) |
| S0 | Scenario 0 | Ambient conditions (specific to this study) |
| S1 | Scenario 1 | Low spring temperatures (specific to this study) |
| S2 | Scenario 2 | Dry spring (specific to this study) |
| S3 | Scenario 3 | Dry previous summer (specific to this study) |
| S4 | Scenario 4 | High autumn temperatures (specific to this study) |
| S5 | Scenario 5 | Slow rising spring temperatures (specific to this study) |
| SLA | Specific Leaf Area | Leaf area divided by its dry mass (measured in mm/mgDW in this study) |
| SOS | Start of growing Season | Point in time at which vegetation begins to reprise its activity after a period of dormancy (typically spring) |
| TRS | Threshold Method | Relative thresholds (of 10%, 20%, 50% in this study). Of the vegetation time series |

**Table 2.**
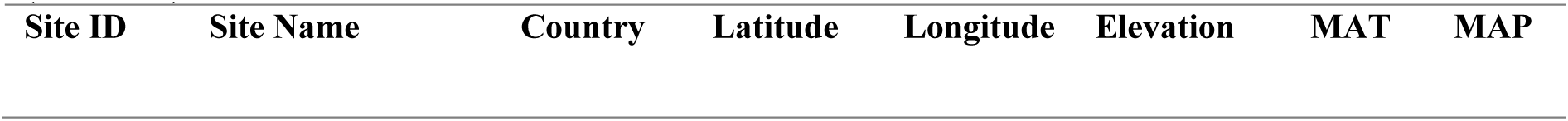
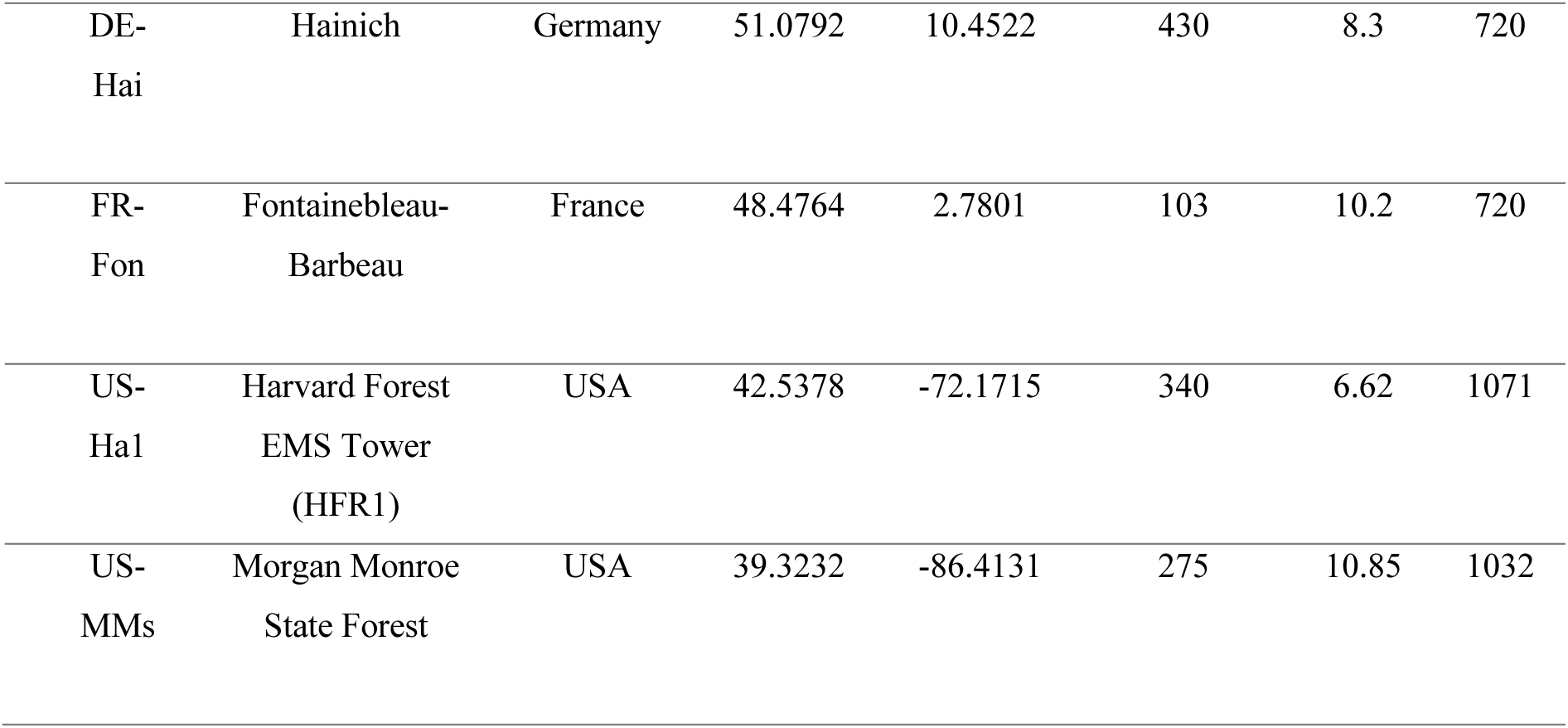
Deciduous broad leaf forest sites selected from the PLUMBER2 dataset, including Site ID, Site Name, Latitude (°), Longitude (°), Elevation (m), Mean Annual Temperature (MAT, °C) and Mean Annual Precipitation (MAP, mm)

### 2.2 Model overview and simulations

For each site, we simulated ecosystem dynamics using QUINCY, a land surface model with fully coupled carbon (C), nitrogen (N), phosphorus (P) and water cycles. QUINCY is structured to follow the flow of these nutrients through soil and vegetation across a set of 14 plant functional types (PFTs) characterised by different leaf type, phenological and photosynthesis strategies. For each PFT, vegetation features structural pools (leaf, fruit, sapwood, heartwood, fine root, coarse root) and storage pools, further divided into labile-where carbon and nutrients are temporarily stored prior to their allocation, and reserve-long term storage. Resources obtained by either photosynthesis or nutrient uptake are thus allocated between pools, balancing between tissue production, maintenance respiration, and the storage of resources for the following season. Resulting plant functions rely on PFT-specific structural characteristics, climatic variables, soil characteristics and nutrients as well as water availability.

Simulated vegetation parameters used in this study are GPP and LAI. QUINCY calculates photosynthesis for sunlit and shaded leaves for each canopy layer following Kull and Kruijt (1998) and photosynthetic capacity depends on leaf N content, which in turn is driven by a balance of N demand for growth and soil available N. Photosynthesis is downregulated in the form of an empirical function by the availability of nutrients and soil moisture, and photosynthetic parameters acclimate to temperature following Friend (2010). Photosynthesis is then integrated to the canopy to derive GPP. The LAI derives from Specific Leaf Area (SLA) characteristic to each PFT (15.4 mm/mgDW for DBF) and leaf mass. The latter depends on the allocation and partitioning of newly assimilated carbon and nutrients to available pools, given by dynamic allometry based on pipe theory and nutrient and water availability. Meristem control further downregulates growth in unfavorable temperature or moisture conditions.

In terms of phenology, meteorological triggers control QUINCY’s start (SOS) and end (EOS) of the growing season, with plant growth set to zero outside of this period. The model differentiates between phenological strategies among PFTs, so that SOS for DBF is a function of the heat accumulation since last dormancy, expressed as growing degree days (*GDD*_acc_) above a PFT-specific temperature threshold (*t_air_^GDD^*), as:

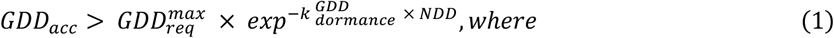

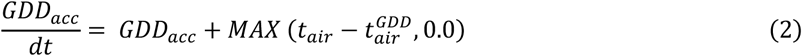

where *GDD*_acc_ is the current growing degree day above *t_air_^GDD^* (5°C for DBF), as the temperature threshold, *t_air_^GDD^* (800.0 °C days for DBF) is a PFT-specific maximum requirement for growing degree days in absence of chilling, −*k_dormance_^GDD^* (0.007 days^-1^ for DBF) is a scaling factor to account for chilling requirements of the buds, *NDD* is the number of days of dormancy since the last growing season, *t*_air_ is the current air temperature (°C), and *dt* is the time-step.

The EOS is in turn triggered by decreasing average air temperatures below a PFT-specific temperature threshold *t_air_^sen^* (8.5 °C for DBF). Temperature-related parameters are expressed in degrees Celsius, *GDD*, *NDD* and *dt* are expressed in days. The SOS and EOS in the model hence indicates the period when plant growth can occur, but the plant growth itself is controlled by environmental conditions as nutrient and water availability, so that LAI and GPP development can be decoupled from the strict growing season even if temperature requirements for SOS are met.

In QUINCY, model simulations are driven using input data for atmospheric conditions, air humidity, soil characteristics, and atmospheric CO_2_ concentrations as well as deposition rates of N, and P. The atmospheric inputs including half-hourly measurements of air temperature, precipitation, incoming radiation, air pressure, humidity, and wind speed, are sourced from site-specific observations provided by the PLUMBER2 dataset (Ukkola et al., 2022). Soil parameters are derived from the SoilGrids database (Hengl, 2017), atmospheric CO_2_ concentrations are based on Le Quéré et al. (2018), nitrogen deposition data come from Lamarque et al. (2010, 2011), and phosphorus deposition is informed by Brahney et al. (2015) and Chien et al. (2016). The model runs with fully transient CO_2_ and nutrient deposition, coupled C-N cycling and prescribed P. Before running the main simulations, QUINCY conducts a spin-up phase of 500 years to stabilize vegetation and soil pools repeating data drawn from 1901–1930, with an accelerator applied to slow soil pools. Following spin-up, simulations for the sites commence in 1901 and continue through 2019.

### 2.3 Phenological scenarios

We conducted simulations under unaltered (S0) and altered meteorological conditions, modifying one climatic forcing variable for each scenario in order to evaluate their individual impact on vegetation processes, and the capacity of each detection method to capture it. Perturbations were imposed to climate data for a year that was common across all sites and did not include extremes (2005). As phenological studies most frequently focus on warm springs (Hassan et al., 2023), we did not include this scenario in our analysis, instead choosing other seasonal perturbation scenarios likely to be caused by climate change. In particular, we tested: (S1) low spring temperatures, with a 2°C temperature reduction from Day of Year (DOY) 32 to 120 (February to April), (S2) spring drought, with a 75% reduction in precipitation from DOY 32 to 151 (February to May), (S3) legacy of summer drought, with a 75% reduction in precipitation from DOY 121 to 212 (May to July) of the previous year, (S4) high autumn temperatures, with a 2°C temperature increase from DOY 213 to 304 (August to October), (S5) slow warming spring, with a 15-day delay in temperature increase from DOY 32 to DOY 120 (February to April).

### 2.4 Calculation of Start and End of Season

From the time series of simulated daily values of GPP and LAI from each scenario, we estimated phenological transition dates using the R package *phenofit* (Kong et al., 2022). To produce a smoothed curve from each time series, we first rough-fitted the data with a weighted Harmonic ANalysis of Time Series (wHANTs) function (Verhoef, 1996; Yang et al., 2015), then performed weight updating using the TIMESAT function (Jönsson & Eklundh, 2004), and last fine fitted the curve using the Elmore logistic approach (Elmore et al., 2012). We selected this combination of functions as the most appropriate to the vegetation type in this study, following recommendations from Kong et al. 2022). Phenological transition dates were then estimated on the resulting smoothed GPP and LAI values for the entire year using:

● *Threshold Method (TRS)*: SOS/EOS are identified as the first days when the normalized vegetation time series crosses set thresholds (10%, 20%, or 50%) of its seasonal amplitude (Fang et al., 2023).
● *Derivative Method (DER)*: SOS/EOS are identified as the days when the first derivative of the vegetation time series (rate of change) reaches its maximum or minimum values (Fisher et al., 2006).
● *Gu Method (GU)*: SOS/EOS are identified as the days where the lower boundaries of the vegetation time series intersect with recovery and senescence lines. These lines are defined by the slopes of two tangent lines that pass through the time series’ maximum and minimum points (Gu et al., 2009).
● *Inflection Method (INFL):* SOS/EOS are identified as the days when the rate of change in the angle of the tangent vector for the vegetation time series reaches a local maximum or minimum (Klosterman et al., 2014; Zhang et al., 2003).

More details for these methods are provided in Kong et al. (2020, 2022).

We then compared the estimated SOS and EOS with the temperature-triggered growing season provided by QUINCY. A workflow overview is presented in Fig.1.

**Figure 1:**
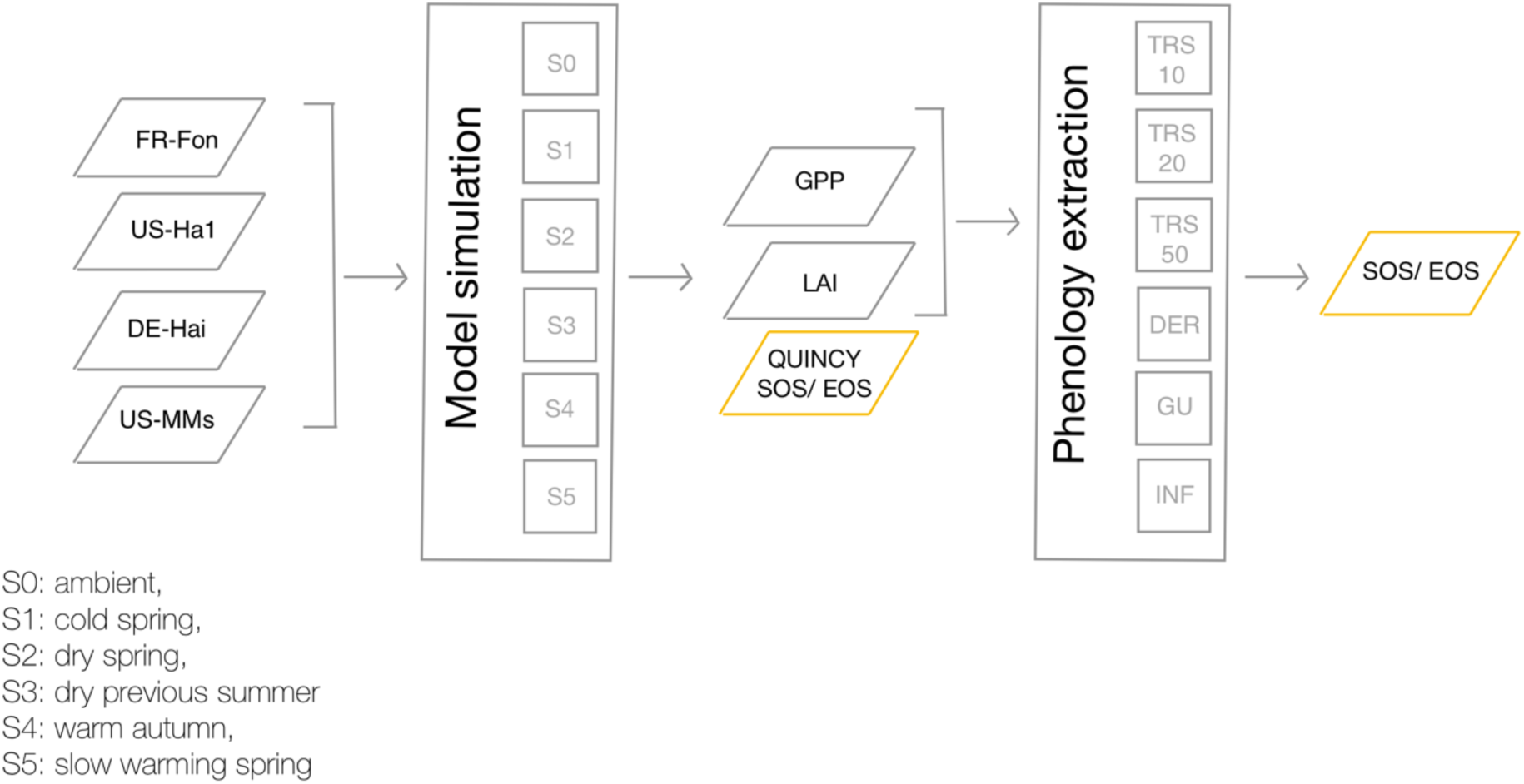
Workflow used for estimating the Start (SOS) and End (EOS) of the growing Season (in yellow) from simulations at four temperate forest sites (FR-Fon, US-Ha1, DE-Hai, US-MMs) across 6 climate scenarios, reported in the figure. QUINCY SOS/ EOS, Gross Primary Productivity (GPP) and Leaf Area Index (LAI) are provided as outputs of each simulation. The latter two are used to extract SOS/ EOS using the workflow from Kong et al. (2022) with Threshold 10% (TRS 10), Threshold 20% (TRS 20), Threshold 50% (TRS 50), Derivative method (DER), Gu method (GU) and Inflection method (INF).

All the analyses were performed in R (version 4.2.2, R Core Team, 2022).

## 3 Results

Under unaltered meteorological conditions, sites exhibit distinct temperature-dependent growing seasons (QUINCY SOS/ EOS in S0, Fig.2-3). Notably, the site FR-Fon displays the longest growing season, as well as the earliest start (234 days, SOS DOY 87, EOS DOY 321), followed by US-MMs (214 days, SOS DOY 108, EOS DOY 322), DE-Hai (194 days, SOS DOY 122, EOS 316), and US-Ha1 (162 days, SOS DOY 134, EOS DOY 296) (Fig. 2-3, Dataset S1). These differences reflect the climatological contexts of each site, where FR-Fon and US-MMs experience the highest temperatures and US-Ha1 on the other hand is characterized by the coldest conditions (Table 2, Fig. S1). Similar patterns are observed in the growing seasons calculated from simulated GPP and LAI values across all six detection methods, with method-specific differences appearing consistently across sites and scenarios (Fig. 2-3, Table S1). Specifically, the inflection method provides the closest dates to the QUINCY SOS across sites and scenarios for both GPP (-2 to 3 days discrepancy) and LAI (up to 4 days discrepancy), although we observe missing EOS dates in US-MMs for S1, S2, S4 and S5. The second closest dates to the QUINCY SOS are provided by the 10% threshold method, with a maximum discrepancy of 4 days for GPP and 7 days for LAI. Notably, the LAI-derived SOS consistently occurs later than GPP SOS, regardless of method or scenario. The closest dates to the QUINCY EOS are instead observed at the 20% threshold for GPP (-9 to 10 days discrepancy), and both the 50% threshold and derivative method for LAI (both showing about 10 days of discrepancy). In general, the 50% threshold and derivative methods delineate the narrowest growing seasons with the latest SOS and earliest EOS and very similar dates, regardless of scenario (Table S1).

**Figure 2.**
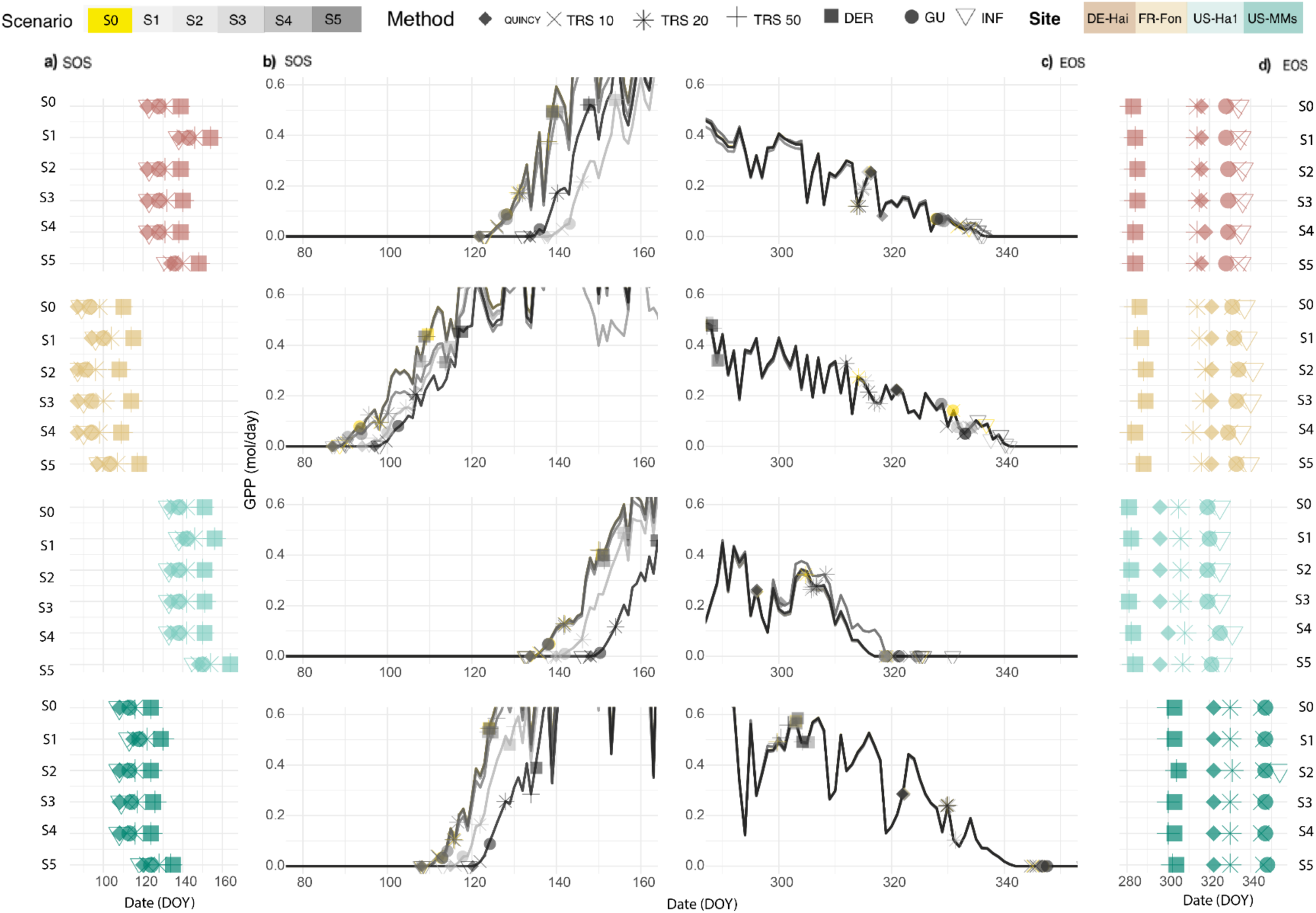
Start (SOS) and End of growing Season (EOS) dates at the sites DE-Hai, FR-Fon, US-Ha1, US-MMs. SOS-EOS are calculated through QUINCY, Threshold 10% (TRS 10), Threshold 20%(TRS 20), Threshold 50% (TRS 50), Derivative method (DER), Gu method (GU) and Inflection method (INF) on simulated GPP. Simulated scenarios are unaltered conditions (S0), cold spring (S1), dry spring (S2), dry previous summer (S3), warm autumn(S4), slow warming spring (S5). In particular (a) shows SOS dates across sites and scenarios, (b) GPP increase across scenarios at SOS and transition dates as provided by methods, (c) GPP decrease across scenarios at EOS and transition dates provided by methods, (d) EOS dates across sites and scenarios.

**Figure 3.**
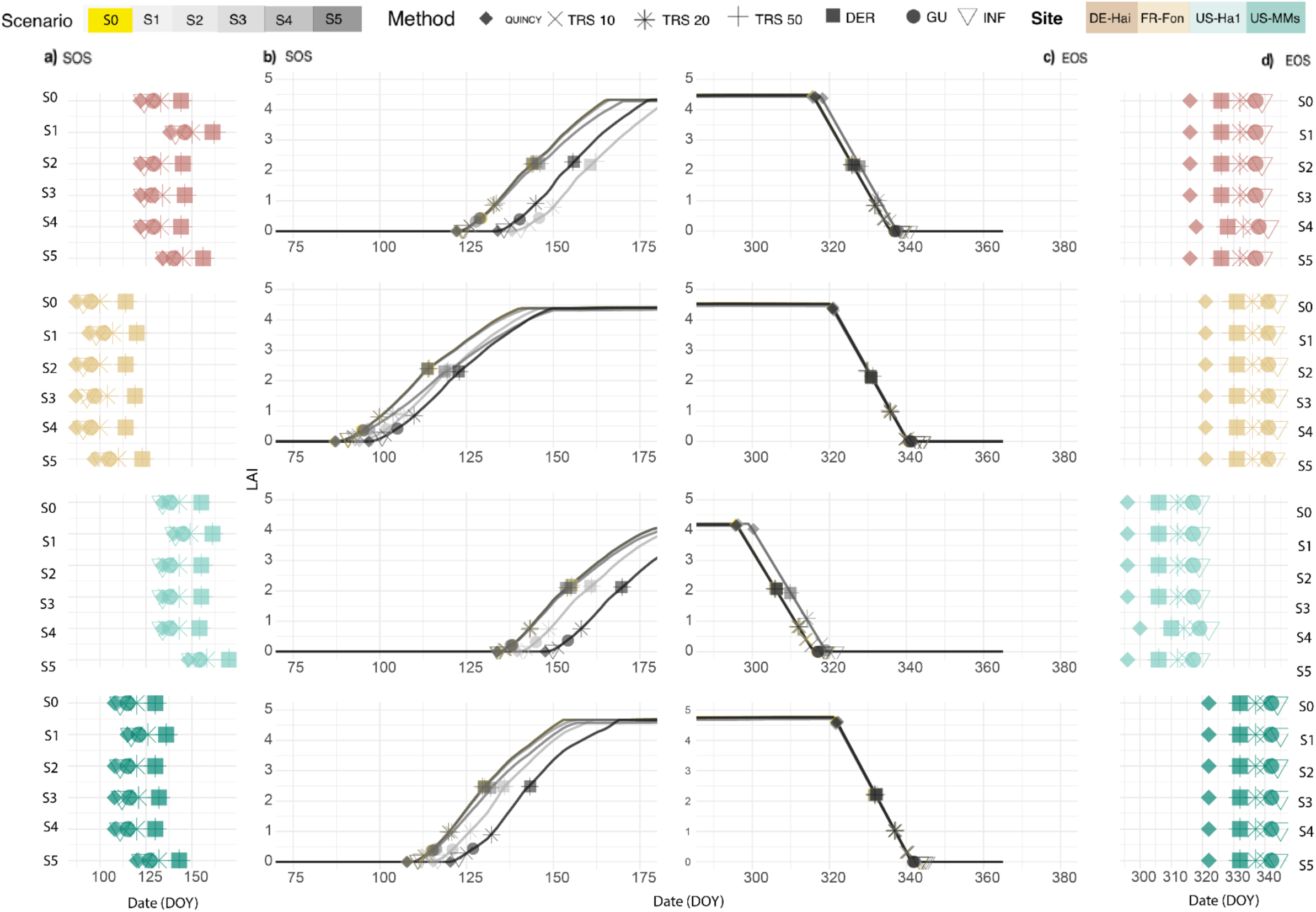
Start (SOS) and End of growing Season (EOS) dates at the sites DE-Hai, FR-Fon, US-Ha1, US-MMs. SOS-EOS are calculated through QUINCY, Threshold 10% (TRS 10), Threshold 20%(TRS 20), Threshold 50% (TRS 50), Derivative method (DER), Gu method (GU) and Inflection method (INF) on simulated LAI. Simulated scenarios are unaltered conditions (S0), cold spring (S1), dry spring (S2), dry previous summer (S3), warm autumn(S4), slow warming spring (S5). In particular (a) shows SOS dates across sites and scenarios, (b) LAI increase across scenarios at SOS and transition dates as provided by methods, (c) LAI decrease across scenarios at EOS and transition dates provided by methods, (d) EOS dates across sites and scenarios

The perturbation scenarios further highlight differences in method sensitivity to site climatic conditions (Fig.4). In the case of low spring temperatures (S1), QUINCY SOS is delayed by 6 to 16 days compared to unaltered conditions, where US-Ha1 is the least and DE-Hai the most affected, with SOS values considerably more delayed in the latter site compared to the others. Similarly, S1 causes a delay in the GPP based SOS regardless of detection method, although to a lesser degree compared to the QUINCY SOS. In particular, the GPP-SOS occurs 1 to 3 days earlier than QUINCY SOS in all sites except for FR-Fon, where the GPP-SOS occurs instead 1 to 2 days later than the QUINCY SOS, depending on method. The LAI-based SOS for S1 are on the other side closer to the QUINCY SOS across detection methods, particularly in FR-Fon and US-Ha1. S1 does not affect QUINCY and LAI EOS, but instead delays GPP EOS of 1 to 2 days, depending on site and detection method.

**Figure 4.**
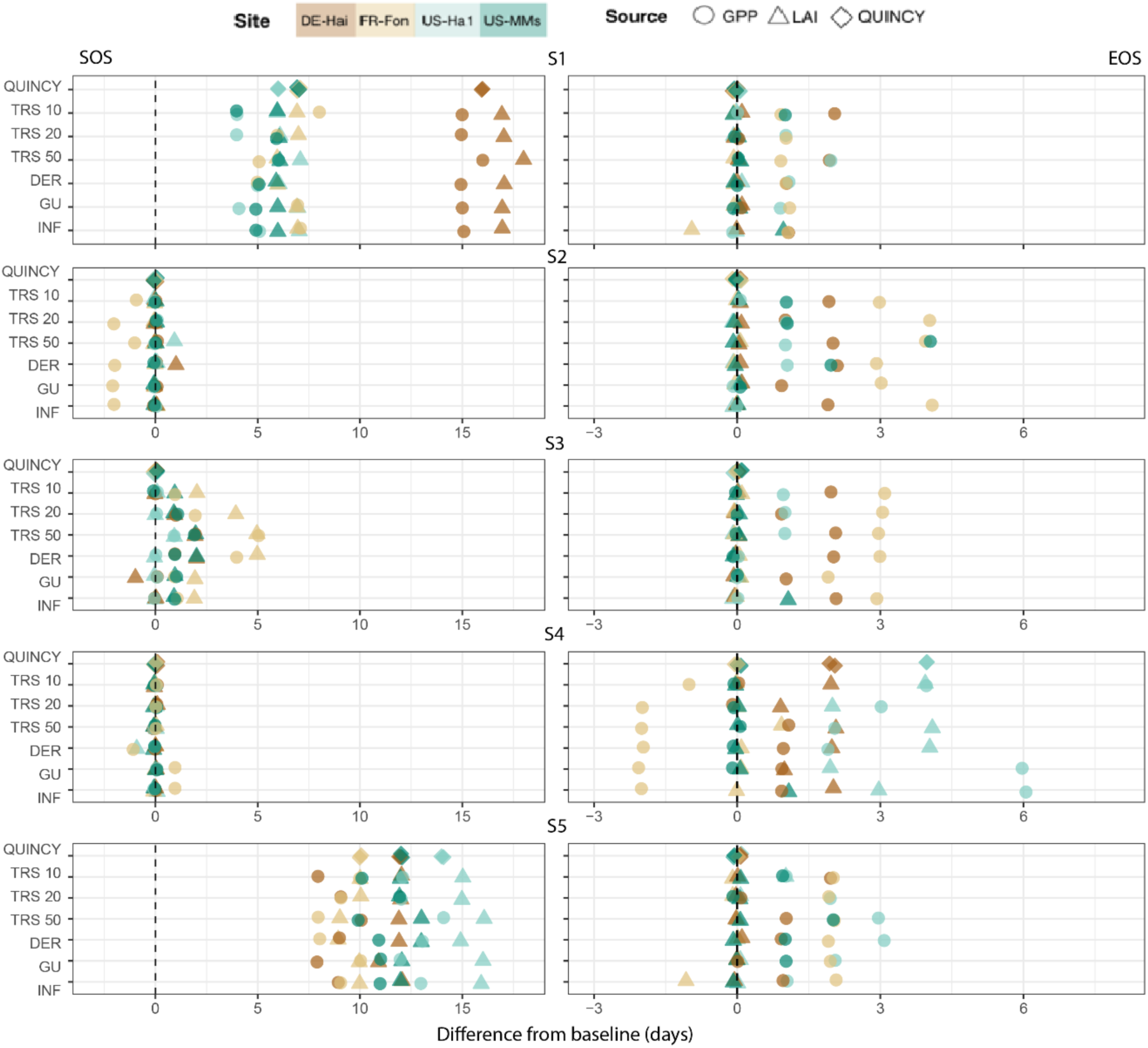
Number of days of difference between Start of Season (SOS) and End of Season (EOS) dates at unaltered conditions (baseline scenario S0, dotted line) and respective dates at cold spring (S1), dry spring (S2), dry previous summer (S3), warm autumn(S4), and slow warming spring (S5) at DE-Hai, FR-Fon, US-Ha1, US-MMs. Dates calculated through QUINCY, Threshold 10% (TRS 10), Threshold 20% (TRS 20), Threshold 50% (TRS 50), Derivative method (DER), Gu method (GU) and Inflection method (INF) on GPP and LAI simulated values

Dry spring conditions (S2) generally do not have any effects on the QUINCY, GPP, and LAI SOS, although with notable exceptions. For instance, an up to 2-day early GPP SOS is evident in FR-Fon across all methods. In the opposite direction, LAI SOS has a 1-day delay in US-Ha1 at the 50% threshold and a 1-day delay in DE-Hai at the derivative method. The GPP EOS also shows a delay of up to 3 days across sites, varying by method.

Summer drought legacy (S3) does not affect the subsequent growing season in QUINCY but impacts both GPP and LAI SOS, showing delays of up to 5 days, depending on site and method. Similar to dry spring conditions, only the GPP EOS shows a delay across sites, with FR-Fon being the most affected site regardless of the method. High autumn temperatures (S4) impact QUINCY EOS only in US-Ha1 and DE-Hai, causing a 4 and 2-days delay respectively. These same sites show similarly delayed GPP and LAI EOS, but in addition to that all methods also detect an up to 2 days early GPP EOS in FR-Fon. Interestingly, FR-Fon also shows a 1-day delay in LAI EOS at the 50% threshold. No effects on SOS were expected as temperature changes have been implemented later in the season, however we still observe 1-day differences in GPP SOS in FR-Fon at the derivative, Gu and Inflection method, and LAI SOS in US-Ha1 at the derivative method.

Under slow warming spring temperatures (S5), the SOS in QUINCY is delayed by 10 to 14 days, again depending on site. The least affected is FR-Fon, while the most pronounced delay occurs in US-Ha1. The GPP SOS is also delayed, although it occurs 1 to 5 days before the corresponding QUINCY SOS, with the largest advance noted in DE-Hai. The LAI SOS is instead generally closer to QUINCY SOS dates, similar to what is observed in S1. Similarly, the EOS for both QUINCY and LAI largely align with minimal variation. Only FR-Fon exhibits a 1-day earlier EOS detected by the inflection method, and US-Ha1 a 1 day late EOS detected by the 10% threshold. In contrast, the GPP EOS shows a consistent delay of 1 to 3 days across all sites and methods

## 4 Discussion

### 4.1 Phenological scenarios

Our work tested the capacity of phenological detection methods to capture under investigated effects of climate change on deciduous broad leaves forests. We intentionally did not address phenological responses to warming conditions at a global scale as they have been widely studied through a variety of methods, including smoothing of time series using singular spectrum analysis (Gu et al., 2022; Keenan et al., 2014), smoothing splines (Keenan et al., 2014; Richardson et al., 2010; Shi et al., 2025), logistic curve fitting (Gu et al., 2022; Keenan et al., 2014; Shi et al., 2025), Savitzky-Golay filter (Gu et al., 2022), asymmetric Gaussian and Harmonic Analysis of Time Series (Shi et al., 2025), and combination of these, but also absence of smoothing (Gu et al., 2022), model prescribed SOS (Ren et al., 2024), and observation of events on the ground (Guralnick et al., 2024; Keenan et al., 2014; Stuble et al., 2021). For time series, dates extraction methods range from relative thresholds (at 10%, 15%, 25%, 30% and 50% depending on the index used) to derivative approaches. A one-to-one comparison of date-extraction methodologies from the literature is complicated by the fact that studies either present outcomes of different methodologies pooled together, or they refer to specific vegetation indexes (e.g. NDVI, GPP, GCC) and time frames. Regardless, all studies agree on a warming induced trend of advanced onset of spring and delayed autumn phenophases, although autumn trends are less consistent (Campioli et al., 2024; Keenan et al., 2014; Stuble et al., 2021), with legacy effects in the following years (Richardson et al., 2010). Such prolonged growing season length increased photosynthetic carbon assimilation, resulting in enhanced carbon uptake (Gu et al., 2022; Ren et al., 2024). This finds an explanation in the influence of high temperatures on photosynthetic enzymatic activity in spring and autumn (Hassan et al., 2023; Richardson et al., 2010) and to decreasing rate of chlorophyll degradation in autumn (Calinger & Curtis, 2023; Campioli et al., 2024; Silvestro et al., 2025). However, phenological responses to future continued warming may still be non-linear (Jochner et al., 2016; Meng et al., 2020), depending on local temperature sensitivity (Fu et al., 2015; Xiong et al., 2024), the velocity of spring time temperature increases (Guralnick et al., 2024), and proximity to vegetation optimal thermal window for growth (Seyednasrollah et al., 2018; Shi et al., 2025).

The influence of local conditions on sites’ responses to climate alterations is also reflected in our simulations, where regardless of date-extraction methodology, GPP, LAI-derived, and QUINCY’s temperature-dependent growing season have the longest duration in warmest sites (FR-Fon, US-MMs), while the coldest and wettest site (US-Ha1) exhibits the latest onset and shortest duration (Fig. 2-3). Regardless of variability between sites, the response to scenarios tested are consistent across methods (Fig. 3). Under cooler or slowly warming spring scenarios (S1, S5), both QUINCY and detection methods show delayed SOS. The EOS is also delayed, but only when derived from GPP, which is more temperature-sensitive due to its direct link to photosynthesis and stress limitations, while LAI shows smoother seasonal changes as its response to temperatures is buffered by its additional dependance on leaf biomass dynamics. In warm autumn conditions (S4), the effects on EOS are manly visible when the latter is derived by GPP, advancing in the driest site (FR-Fon) but is being in colder sites (US-Ha1, DE-Hai). Non-temperature related scenarios primarily affect GPP and LAI-derived seasons as spring drought (S2) advances GPP SOS in FR-Fon, while summer drought legacy (S3) delays both GPP and LAI SOS but only prolongs GPP seasons, likely due to reduced growth and storage in the prior season.

In general simulations reflect the expected behavior of temperate forest phenology and are consistent with the literature (Q. Wang et al., 2024; Xiong et al., 2023), with differential effects between sites reflecting the influence of local conditions (Bórnez et al., 2021; Duan et al., 2018, Q. Wang et al., 2024). Different sensitivity of simulated processes to environmental changes is further discussed in section 4.3.

### 4.2 Phenological detection methods and specificities

Between-method differences in phenological dates are maintained across scenarios. The inflection method provides the closest SOS dates to QUINCY for both GPP and LAI, as it detects local maxima/minima in the curvature’s rate of change (Zhang et al., 2003), aligning with ground-based, temperature-dependent leaf-out dates (Klosterman et al., 2014). However, its reliance on four inflection points can lead to missing data when conditions are not met (Kong et al., 2022), as seen with missing GPP EOS dates in US-MMs (Table S1, Dataset S1), likely due to high temperatures and precipitation (Figure 2). The 10% threshold is the second closest to QUINCY SOS, reflecting early plant activity (Panwar et al., 2023). For EOS, most methods yield dates later than QUINCY, especially for LAI (Fig. 3, Table S1), attributed to model parametrization and the timing of LAI decrease as leaf shedding only starts in the model once the EOS trigger is reached. The closest EOS dates to QUINCY are from the 20% threshold for GPP (Fig. 2) and 50% threshold/derivative method for LAI (Fig. 3), supporting the use of lower thresholds for SOS and higher for EOS in the context of this study (Panwar et al., 2023). The 50% threshold and derivative method consistently show the latest SOS and earliest EOS, defining the narrowest growing season and demarcating the peak of season (POS). The reliance of POS dynamics on a different set of factors other than SOS/ EOS such as early-season vegetation activity and developmental constraints, and the effect of POS shifts on maximum resource availability support the need to study it as separate from SOS/ EOS dynamics, a point already noted in the literature (Li et al., 2025; Liu et al., 2025). These methods indeed respond distinctly to simulated conditions, particularly under summer drought legacies (S3), with pronounced delays in SOS at the driest site, FR-Fon.

It is worth highlighting that while phenological dates are generally consistent across methods, some exceptions occur. For instance, summer drought (S3) in DE-Hai delays SOS using the 50% threshold and derivative methods, yet advances it by one day with the Gu method. Another case can be made for the warm autumn scenario (S4) showing one-day shifts in SOS under the derivative, Gu, and inflection methods, despite changes in conditions occurring later in the season. These discrepancies, rarely exceeding one day, likely arise from SOS/EOS calculations being based on annual fitted values, reflecting year-round rather than seasonal variations. Detection methods depend on the shape of the vegetation time series, so any factor altering this shape can impact phenological assessments.

### 4.3 Beyond temperature threshold effects in models

This study has shown that, for the scenarios investigated, non-temperature driven changes in climatic conditions have effects on the growing season, including between-site dynamics, which would not be picked up by temperature threshold methods and models but can still be captured by standard methods for detecting SOS and EOS. In particular, we observe SOS-POS-EOS responses to environmental changes detected on simulated GPP and LAI that are missed in the QUINCY growing season (Fig.3), where internal SOS/ EOS are triggered solely by temperature, a common approach in phenology models (Dávid et al., 2021; Fu et al., 2014; Mo et al., 2023a). Phenological models that exclusively rely on temperatures oversee factors that influence plant responses to environmental changes, such as the joint effects of chilling, forcing, photoperiod, precipitation (Liang et al., 2024; Mo, et al., 2023b; Mo, Zhang, et al., 2023), and legacy effects (Keenan & Richardson, 2015; Liu et al., 2024), and ultimately affect the accuracy of the simulated growing season. Further, this effect might be missed in regional or global studies that correlate phenological dates derived from vegetation time series with increases in temperatures, without taking into account other drivers (Dávid et al., 2021b; J. Wang et al., 2021). A more realistic representation of the growing season beyond GDD type thresholds in LSMs (e.g. Caldararu et al., 2012, 2014, Seitz et al., 2026) would be necessary for accounting said dynamics and better simulate plant phenology (Mo, et al., 2023b).

### 4.4 Models as a tool for theoretical exploration

Process based models such as QUINCY create simulated data that are ecologically realistic and that can easily be manipulated, essentially providing a potential large number of ecosystem experiments *in silico*. The generation of synthetic data from models for testing is a known approach in ecology (Zurell et al., 2010) and found its application in various fields (DiRenzo et al., 2023; Fernández et al., 2024; Figueira & Vaz, 2022; Rahman et al., 2020; Zhan et al., 2022). LSMs however are still predominantly treated as predictive tools, where ecological data provide the ‘true’ ground for evaluation. It must be acknowledged that either LSMs or land components of Earth System Models have been used as controlled experiments to investigate the impact of specific process shifts on the land–atmosphere coupling (Li et al., 2023, 2024; Zhu et al., 2022); however, this approach can still be considered bound to forecasting objectives. Here we use QUINCY simulations as baseline ‘truth’-we know when the growing season starts and ends and we can alter environmental conditions to explore the agreement between phenological detection methods. A similar philosophy is shared by Observing System Simulation Experiments (OSSEs), which generate synthetic observations drawn from a nature run, i.e. a baseline perturbation-free run from atmospheric or Earth System Models, against which outputs from other runs are compared (Hoffman & Atlas, 2016).

However, OSSEs are specifically tailored to assess data assimilation pipelines and instrument configurations, thus their application has remained limited to the evaluation of new or proposed observation systems (Liu et al., 2021; Pflug et al., 2024; Rahman et al., 2020; Wang et al., 2023). Regardless of the approach, it remains of course important to keep in mind the limitations of using model simulations, as any assumptions already existing in the model is incorporated in the study. For example in the context of this study, the GDD and chilling requirement parameterisation affects the magnitude of changes in any of our scenarios and implicitly the capacity of each method to detect such changes. Representations of soil hydrology and plant response to water stress will in turn influence the response to the water-based scenarios. Future studies could include multiple LSMs and a more comprehensive uncertainty analysis.

## 5 Conclusion

In this study we compare commonly used phenological detection methodologies on four temperate forest sites, where we use the QUINCY LSM to simulate GPP and LAI under five different climate scenarios. Despite method specific differences that remain largely consistent across scenarios, the growing season provided by model simulations and methods largely agree, responding to changes according to site-specific environmental context. Methods sensitive to the earliest alterations provide the closest match to a temperature-only dependent SOS, while higher thresholds are closer to the EOS provided by the model. This likely reflects the reliance of early spring and autumn phenology events on temperatures. Highest thresholds and methods based on the rate of change of vegetation parameters delimit peak of season, which can be affected differentially by environmental forcings. We also show how some methods, despite broadly agreeing in their response to simulated changes, are also able to pick up on effects that were overlooked by the GDD-threshold type growing season, warranting discussion in favor of a more realistic representations of phenology in process-based models. Despite these limitations, QUINCY provided good ground for the generation of synthetic data to be used for testing phenological detection methodologies, supporting the use of LSMs beyond their current predictive application into tools for theoretical exploration.

## Code and data availability

The data that support the findings of this study is available in the Supporting Information. The R code used to generate the data is permanently archived in Zenodo at https://doi.org/10.5281/zenodo.21904407. The input PLUMBER2 NetCDF data are publicly available at https://dx.doi.org/10.25914/5fdb0902607e1. The QUINCY model’s source-code is available at https://git.bgc-jena.mpg.de/quincy-model/qs-open-access. More information at https://doi.org/10.17871/quincy-model-2019.

## Author contributions

SC conceptualised the study and supervised the work, MY conducted the analyses, JS tested the workflow for extracting phenological dates, WJM, LD and GGN provided feedbacks on the data processing. All authors contributed to the revision of the manuscript.

## Statement on inclusion in global research

This study is based entirely on simulated data. All empirical data used to inform the simulations were obtained from publicly available repositories, cited in the text. Consequently, no local data were collected. The simulated sites represent ecosystems in the Northern Hemisphere, contributing to the existing body of ecological research focused on this region. While this focus reflects the availability of data and established modelling frameworks, we acknowledge that ecological patterns and processes may differ in other underrepresented regions.

## Disclaimer

Copernicus Publications remains neutral with regard to jurisdictional claims made in the text, published maps, institutional affiliations, or any other geographical representation in this paper. While Copernicus Publications makes every effort to include appropriate place names, the final responsibility lies with the authors. Views expressed in the text are those of the authors and do not necessarily reflect the views of the publisher.

## Acknowledgements

We thank Dr Camille Abadie for her thoughtful feedback and helpful suggestions, which contributed to the improvement of this manuscript. We acknowledge the use of Google Gemini (version 3.6 Flash, Google, https://gemini.google.com) to refine the grammar and flow of sections of the text. All AI-generated outputs were reviewed, verified, and modified by the corresponding author. This work was supported by RIT (Research IT, Trinity College Dublin). All calculations were performed on the Callan cluster maintained by the Trinity Centre for High Performance Computing.

