## Appendix for "Benchmarking phenology with a land surface model: an in- silico experiment for temperate forests"

### Appendix B: Figures

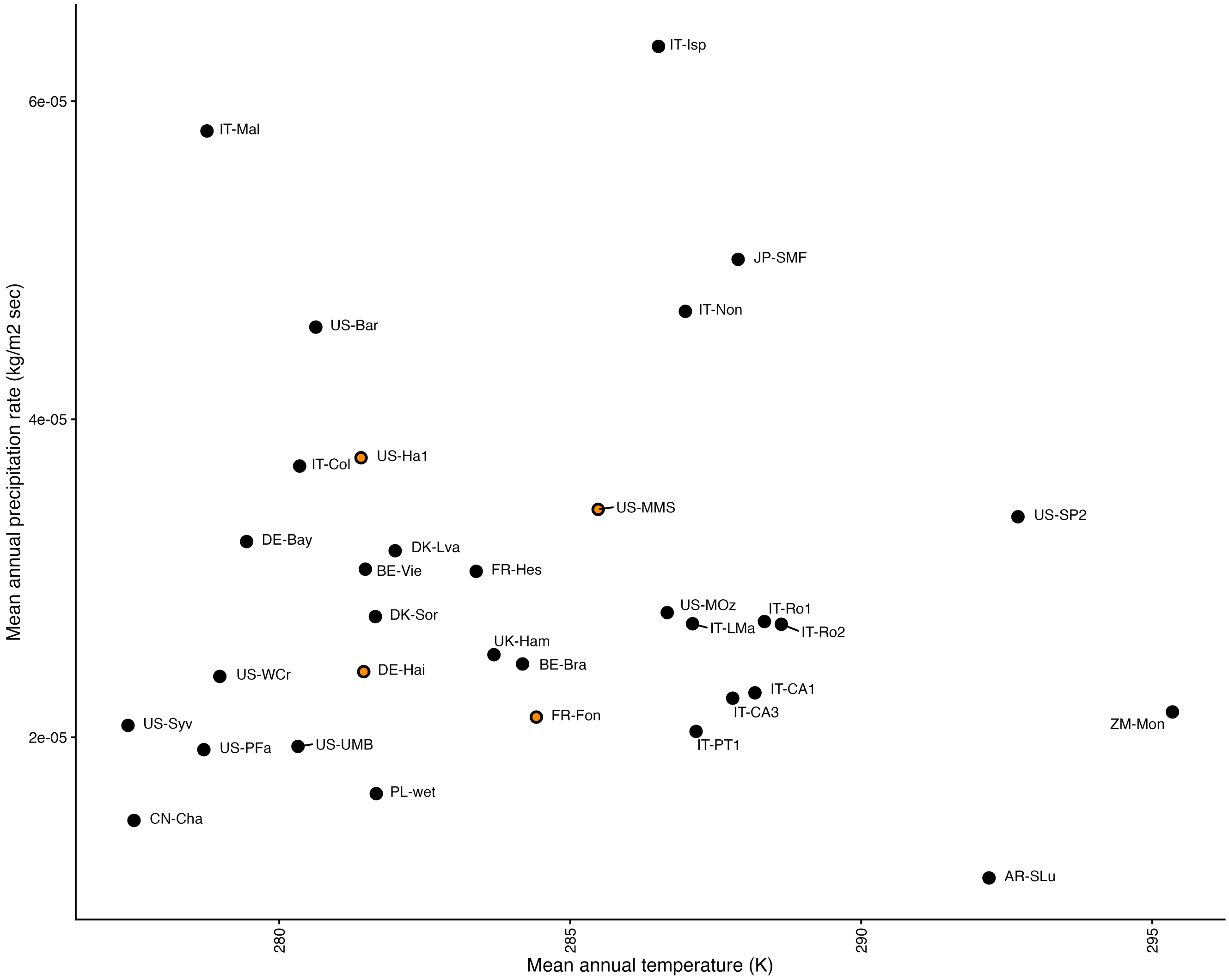


Figure B1: Mean annual precipitation rate (kg/m2 sec) and mean annual temperature (K) for PLUMBER2 deciduous broad-leaf forest sites

### Appendix C: Tables

| Source | Site | Scenario | Sos TRS 10% | Sos TRS 20% | Sos TRS 50% | Sos DER | Sos Gu | Sos Inflection | Eos TRS 10% | Eos TRS 20% | Eos TRS 50% | Eos DER | Eos Gu | Eos Inflection |
|---|---|---|---|---|---|---|---|---|---|---|---|---|---|---|
| GPP | DE-Hai | 0 | 4 | 9 | 16 | 17 | 6 | 1 | 16 | -2 | -34 | -33 | 12 | 18 |
| GPP | DE-Hai | 1 | 3 | 8 | 16 | 16 | 5 | 0 | 18 | -2 | -32 | -32 | 12 | 19 |
| GPP | DE-Hai | 2 | 4 | 9 | 16 | 17 | 6 | 1 | 18 | -1 | -32 | -31 | 13 | 20 |
| GPP | DE-Hai | 3 | 4 | 10 | 18 | 18 | 6 | 1 | 18 | -1 | -32 | -31 | 13 | 20 |
| GPP | DE-Hai | 4 | 4 | 9 | 16 | 17 | 6 | 1 | 14 | -4 | -35 | -34 | 11 | 17 |
| GPP | DE-Hai | 5 | 0 | 6 | 14 | 14 | 2 | -2 | 18 | -2 | -33 | -32 | 12 | 19 |
| GPP | FR-Fon | 0 | 3 | 11 | 22 | 23 | 6 | 2 | 10 | -7 | -37 | -35 | 10 | 16 |
| GPP | FR-Fon | 1 | 4 | 10 | 20 | 21 | 6 | 2 | 11 | -6 | -36 | -34 | 11 | 17 |
| GPP | FR-Fon | 2 | 2 | 9 | 21 | 21 | 4 | 0 | 13 | -3 | -33 | -32 | 13 | 20 |
| GPP | FR-Fon | 3 | 4 | 13 | 27 | 27 | 7 | 3 | 13 | -4 | -34 | -32 | 12 | 19 |
| GPP | FR-Fon | 4 | 3 | 11 | 22 | 22 | 7 | 3 | 9 | -9 | -39 | -37 | 8 | 14 |
| GPP | FR-Fon | 5 | 3 | 10 | 20 | 21 | 6 | 1 | 12 | -5 | -35 | -33 | 12 | 18 |
| GPP | US-Ha1 | 0 | 2 | 8 | 16 | 17 | 4 | -1 | 23 | 8 | -18 | -17 | 25 | 31 |
| GPP | US-Ha1 | 1 | 0 | 6 | 16 | 16 | 2 | -2 | 23 | 10 | -14 | -14 | 24 | 29 |
| GPP | US-Ha1 | 2 | 2 | 8 | 16 | 17 | 4 | -1 | 23 | 9 | -16 | -15 | 23 | 29 |
| GPP | US-Ha1 | 3 | 0 | 6 | 16 | 16 | 2 | -2 | 24 | 11 | -13 | -12 | 25 | 30 |
| GPP | US-Ha1 | 4 | 2 | 8 | 17 | 17 | 4 | -1 | 24 | 10 | -15 | -15 | 23 | 29 |
| GPP | US-Ha1 | 5 | 2 | 8 | 16 | 17 | 4 | -1 | 23 | 10 | -15 | -14 | 23 | 29 |
| GPP | US-MMs | 0 | 2 | 8 | 14 | 15 | 4 | -1 | 24 | 8 | -20 | -18 | 26 | NA |
| GPP | US-MMs | 1 | 4 | 8 | 16 | 16 | 5 | 0 | 23 | 8 | -22 | -19 | 25 | NA |
| GPP | US-MMs | 2 | 4 | 9 | 18 | 17 | 6 | 1 | 23 | 8 | -22 | -19 | 25 | NA |

| | | | | | | | | | | | | | | |
|---|---|---|---|---|---|---|---|---|---|---|---|---|---|---|
| GPP | US-MMs | 3 | 4 | 8 | 16 | 16 | 5 | 0 | 24 | 9 | -18 | -17 | 25 | 32 |
| GPP | US-MMs | 4 | 1 | 7 | 15 | 14 | 3 | -2 | 24 | 8 | -22 | -19 | 25 | NA |
| GPP | US-MMs | 5 | 4 | 8 | 16 | 16 | 5 | 0 | 23 | 8 | -22 | -19 | 25 | NA |
| LAI | DE-Hai | 0 | 4 | 11 | 22 | 22 | 7 | 2 | 18 | 16 | 10 | 10 | 21 | 23 |
| LAI | DE-Hai | 1 | 5 | 12 | 24 | 23 | 8 | 3 | 18 | 16 | 10 | 10 | 21 | 23 |
| LAI | DE-Hai | 2 | 4 | 11 | 22 | 23 | 7 | 2 | 18 | 16 | 10 | 10 | 21 | 23 |
| LAI | DE-Hai | 3 | 4 | 12 | 24 | 24 | 6 | 2 | 18 | 16 | 10 | 10 | 21 | 23 |
| LAI | DE-Hai | 4 | 4 | 11 | 22 | 22 | 7 | 2 | 18 | 15 | 10 | 10 | 20 | 23 |
| LAI | DE-Hai | 5 | 4 | 11 | 22 | 22 | 6 | 2 | 18 | 16 | 10 | 10 | 21 | 23 |
| LAI | FR-Fon | 0 | 5 | 13 | 27 | 27 | 8 | 4 | 19 | 15 | 9 | 10 | 20 | 23 |
| LAI | FR-Fon | 1 | 5 | 13 | 26 | 26 | 8 | 4 | 19 | 15 | 9 | 10 | 20 | 22 |
| LAI | FR-Fon | 2 | 5 | 13 | 27 | 27 | 8 | 4 | 19 | 15 | 9 | 10 | 20 | 23 |
| LAI | FR-Fon | 3 | 7 | 17 | 32 | 32 | 10 | 6 | 19 | 15 | 9 | 10 | 20 | 23 |
| LAI | FR-Fon | 4 | 5 | 13 | 27 | 27 | 8 | 4 | 19 | 15 | 10 | 10 | 20 | 23 |
| LAI | FR-Fon | 5 | 5 | 13 | 26 | 26 | 8 | 4 | 19 | 15 | 9 | 10 | 20 | 22 |
| LAI | US-Ha1 | 0 | 2 | 9 | 20 | 21 | 4 | 0 | 18 | 16 | 10 | 10 | 21 | 23 |
| LAI | US-Ha1 | 1 | 2 | 9 | 21 | 21 | 5 | 1 | 18 | 16 | 10 | 10 | 21 | 23 |
| LAI | US-Ha1 | 2 | 2 | 9 | 21 | 21 | 4 | 0 | 18 | 16 | 10 | 10 | 21 | 23 |
| LAI | US-Ha1 | 3 | 2 | 9 | 21 | 21 | 4 | 0 | 18 | 16 | 10 | 10 | 21 | 23 |
| LAI | US-Ha1 | 4 | 2 | 9 | 20 | 20 | 4 | 0 | 18 | 14 | 10 | 10 | 19 | 22 |
| LAI | US-Ha1 | 5 | 3 | 10 | 22 | 22 | 6 | 2 | 19 | 16 | 10 | 10 | 21 | 23 |
| LAI | US-MMs | 0 | 5 | 12 | 22 | 22 | 7 | 3 | 18 | 15 | 10 | 10 | 20 | 22 |
| LAI | US-MMs | 1 | 4 | 11 | 21 | 21 | 6 | 2 | 18 | 15 | 10 | 10 | 20 | 23 |
| LAI | US-MMs | 2 | 5 | 12 | 22 | 22 | 7 | 3 | 18 | 15 | 10 | 10 | 20 | 22 |
| LAI | US-MMs | 3 | 6 | 13 | 24 | 24 | 8 | 4 | 18 | 15 | 10 | 10 | 20 | 23 |

| | | | | | | | | | | | | | | |
|---|---|---|---|---|---|---|---|---|---|---|---|---|---|---|
| LAI | US-MMs | 4 | 5 | 12 | 22 | 22 | 7 | 3 | 18 | 15 | 10 | 10 | 20 | 23 |
| LAI | US-MMs | 5 | 5 | 12 | 23 | 23 | 7 | 3 | 18 | 15 | 10 | 10 | 20 | 22 |

**Table C1:** Number of days of difference between QUINCY and methods-derived SOS/EOS for each site and scenario. Table is color-coded to highlight closest dates to QUINCY SOS (white – closest, green - furthest) and EOS (white – closet, brown - furthest)
